# Transcriptomic profiling of plaque psoriasis and cutaneous T cell subsets during treatment with secukinumab

**DOI:** 10.1101/764118

**Authors:** Jared Liu, Hsin-Wen Chang, Kristen M. Beck, Sahil Sekhon, Timothy H. Schmidt, Di Yan, Zhi-Ming Huang, Eric J. Yang, Isabelle M. Sanchez, Mio Nakamura, Shrishti Bhattarai, Quinn Thibodeaux, Richard Ahn, Tina Bhutani, Michael D. Rosenblum, Wilson Liao

## Abstract

The IL17A inhibitor secukinumab is efficacious for the treatment of psoriasis. In order to define its mechanism of action, it is important to understand its impact on psoriatic whole skin tissue as well as specific skin-resident immune cell populations such as T lymphocytes. In this study, we treated 15 moderate-to-severe plaque psoriasis patients with secukinumab and characterized the longitudinal transcriptomic changes of whole lesional skin tissue and cutaneous CD4^+^ T effector cells (Teffs), CD4^+^ T regulatory cells (Tregs), and CD8^+^ T effector cells during 12 weeks of treatment. Secukinumab was clinically effective, with 100%, 47%, and 27% of patients in the study achieving PASI75, PASI90, and PASI100 by week 12, respectively. At baseline prior to treatment, we observed that *IL17A* overexpression predominates in psoriatic CD8^+^ T cells rather than Teffs, supporting the importance of IL-17-secreting CD8+ T cells (Tc17) compared to IL-17-secreting CD4+ T cells (Th17) cells in the pathogenesis of psoriasis. Although secukinumab targets only *IL17A*, we observed rapid reduction of *IL17A, IL17F, IL23A, IL23R*, and *IFNG* expression in lesional skin as soon as 2 weeks after initiation of treatment and normalization of expression by week 12. Secukinumab treatment resulted in resolution of 89-97% of psoriasis-associated expression differences in both bulk tissue and T cell subsets by week 12 of treatment. Overall, secukinumab appears to rapidly reverse many of the molecular hallmarks of psoriasis.

## Introduction

Psoriasis is a chronic inflammatory disease of the skin affecting about 2% of the population [1]. This disease primarily manifests as scaly, epidermal plaques formed by a hyperproliferation of abnormally differentiating keratinocytes and an infiltration of activated innate and adaptive immune cells. These pathological changes are currently understood to result from an immune-mediated response to autoantigens whose diversity and pathogenic origins are still largely unknown.

The cytokine IL-17A has been found to drive much of the cellular pathology of psoriasis. It is produced by a number of infiltrating immune cells types including T cells, neutrophils, and mast cells in psoriatic lesions [2–4], where it induces proliferation, hinders differentiation, and upregulates the expression of inflammatory cytokines in keratinocytes [5,6], thereby promoting and maintaining the characteristic physiopathology of psoriasis.

Supporting a central role for IL-17A in psoriasis is the success of the IL-17A blocker secukinumab in the treatment of this disease. Secukinumab was found in two major phase 3 clinical trials to be efficacious in the treatment of psoriasis, achieving a PASI75 response in over 70% of psoriasis patients, higher than the response rate achieved by the TNF blocker etanercept [7]. At the transcriptomic level, clinical improvement in response to secukinumab was found to correlate with a resolution of disease-associated gene expression (i.e. genes differentially expressed between pre-treatment psoriatic skin and healthy skin) in lesional tissue, notably in genes involved in Th17 signaling and development, such as *IL17A*, *IL23A*, and *IFNG* [8,9]. Presently, it is still unclear how secukinumab affects specific resident cell types in the lesion, particularly T cells, which are currently understood to play an important role in initiating psoriasis pathogenesis in the context of IL-17A signaling. Th17 cells occur at higher frequency in psoriatic lesions compared with healthy skin [4], and several identified psoriasis autoantigens, such as LL37 [10] and the melanocyte protein ADAMTSL5 [11], have been found to activate CD4^+^ and CD8^+^ T cells to produce IL-17A.

In this study, we use RNA-seq to characterize the transcriptomic phenotype of T cell subsets along with whole tissue within psoriatic lesions as well as their response to secukinumab treatment, focusing on the genes and pathways altered by disease and by treatment.

## Methods

### Patients

This study was conducted at a single academic center. Patients were at least 18 years of age with a diagnosis of moderate to severe plaque psoriasis for at least 6 months or longer. Moderate to severe psoriasis was defined as a score of 12 or higher on the Psoriasis Area and Severity Index (PASI), and a score of 3 or higher on the Physician’s Global Assessment (PGA). Patients could have received previous systemic treatment or phototherapy, including anti-TNF, anti-IL-12/23, and anti-IL-17 biologic agents; however, a washout period of 2 weeks for topicals and phototherapy, 4 weeks for oral systemic medications, 3 months for most biologics, and 6 months for ustekinumab were required. Study subjects received secukinumab (300 mg, subcutaneous) at weeks 0, 1, 2, 3, and 4, then every four weeks thereafter until week 48. Skin punch biopsies were obtained at weeks 0, 2, 4, and 12. Biopsies were taken from the edge of a psoriatic plaque, and subsequent biopsies in a given patient were taken from the same resolving plaque or an anatomically similar plaque. PASI and PGA assessments were performed at weeks 0, 4, 8, 12, 16, 24, 36, and 52.

### Cell Sorting

Skin punch biopsies were immediately stored at 4 °C in a container with sterile gauze and PBS until they were ready to be processed. The tissue was trimmed to remove hair and subcutaneous adipose before being finely minced and mixed with digestion buffer composed of Collagenase Type IV (Worthington LS04188), DNAse (Sigma-Aldrich DN25-1G), 10% FBS, 1% HEPES, and 1% Penicillin/Streptavidin in RPMI 1640. After overnight incubation in 5% CO2, cell suspensions were harvested, filtered, centrifuged, counted and sorted for RNA-seq experiments. For cell sorting, a lymphocyte gate was taken and then doublets were excluded in a following gate. From there, live CD45^+^ cells were gated, then CD45^+^ CD3^+^ cells, and from here CD4^+^ and CD8^+^ cells could be observed. From the CD4^+^ gate, CD25 was plotted vs CD27. CD25^−^ CD27^−^ cells were gated and called T effectors, and CD25^+^CD27^+^ cells were called Tregs. The cells were sorted into 1X lysis buffer from the SMART-Seq v4 Ultra Low Input RNA Kit (Takara Clontech cat. 634894) and snap frozen in liquid nitrogen for storage at −80 °C. Reported post-sort purity was greater than 90% for all populations.

### Sample preparation for RNA-Seq

Sequencing libraries were constructed with sorted lymphocyte lysates and purified RNA extracted from whole skin tissue using the SMART-Seq v4 Ultra Low Input RNA Kit (Takara Clontech cat. 634894). Bulk RNA was extracted from whole skin tissue using Qiagen RNeasy fibrous tissue mini kit (cat. 74704). Reverse transcription was performed on poly-adenylated RNA from cell lysate for up to 1000 cells followed by second strand synthesis to make full length cDNA. cDNA was then later amplified by PCR using cycle numbers determined by cell numbers. Amplified cDNA was purified by AMPure XP beads (Beckman Coulter) and followed by Qubit quantification. Sequencing libraries were constructed using the Illumina Nextera DNA kit with 0.4 ng of amplified cDNA for sorted lymphocytes and 0.3ng for cDNA synthesized from bulk skin RNA. The sizes of constructed libraries were evaluated by Illumina Bioanalyzer High Sensitivity DNA Assay to ensure library sizes range between 300bp to 500bp. The libraries were quantified using NEBNext Library Quant Kit for Illumina (NEB #E7630) to ensure equal molar concentrations of sequenceable libraries were pooled. Sequencing reactions were performed on Illumina Hiseq4000 with pair-end 150 bp reads. More than 30 million reads were generated per sample.

### RNA-seq QC and data processing

Reads were mapped to the human genome (GRCh38, with Gencode v25 annotation) using STAR v2.4.2a [12]. Picard-tools v1.139 (http://broadinstitute.github.io/picard) was used to compile mapping statistics for the resulting alignments, and a table of counts of each gene from each sample was generated with HTSeq v0.6.1p1 [13]. Samples with less than 1 million reads, 10,000 detected genes, or 60% mappable reads were excluded. Genes expressed at fewer than 5 reads in fewer than 20% of samples for each cell type were also excluded from further analysis. We used DESeq2 v1.18.1 [14] to normalize read counts for each gene between samples and to generate linear models for differential expression analysis, accounting for technical effects and biological variation using the following formula for bulk tissue:

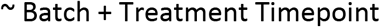

For sorted cells, we accounted for additional technical effects using the following formula:

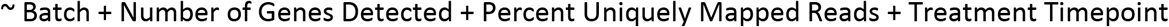

Genes differentially expressed between groups were calculated with DESeq2 using the Wald test with parametric fit. Principal components analysis (PCA) was performed using the DESeq2 function ‘plotPCA’ on variance-stabilized count data (using the DEseq2 function ‘varianceStabilizingTransformation’) that was adjusted for the modeled effects of batch, number of genes detected, and percent uniquely mapped reads for all sample types.

### Pathway analysis

We used the Panther database[15] to identify overrepresented biological processes and pathways from lists of differentially expressed genes, based on Fisher’s exact test with p-values corrected by FDR. Mann-Whitney rank-sum test was used to determine if the log fold changes of genes associated to a biological process or pathway are enriched compared to the overall distribution of the whole gene list.

## Results

We characterized the clinical and molecular responses of 15 psoriasis patients who received secukinumab (300 mg) over the course of the 52-week study interval (Figure 1A). The patient demographics are summarized in Table 1. A total of 60% of the patients were male, the median age was 37 years, and the mean duration of psoriasis was approximately 15 years. The mean PASI score was 20.8, 80% of the patients had a baseline PGA score of 4 or 5, and 87% had previously been treated with either phototherapy, conventional systemic therapy, or biologic therapy. Additional patient information is included in Supplementary Table 1. Of the total 15 patients enrolled, 13 patients completed all 52 weeks of the study, as one patient was lost to follow up after week 36 and another was discontinued from the study due to a neurologic event after week 24, which was thought to be unrelated to the study treatment.

**Table 1.** Patient Characteristics*

| Characteristic | Patients (N=15) |
| --- | --- |
| Median age, years | 37 |
| Male sex, no. (%) | 9 (60%) |
| Race/Ethnicity, no. (%)^ |  |
| Asian/Pacific Islander | 4 (27%) |
| Black/African American | 2 (13%) |
| Caucasian | 5 (33%) |
| Hispanic | 4 (27%) |
| Duration of psoriasis, years | 15 (±10) |
| PASI score | 20.8±13.4 |
| PGA score |  |
| 3 | 3 (20%) |
| 4 | 9 (60%) |
| 5 | 3 (20%) |
| Psoriatic arthritis, no. (%) | 1 (7%) |
| Family history of psoriasis, no. (%) | 5 (33%) |
| Previous treatment, no. (%) |  |
| Topical therapy only | 2 (13%) |
| Phototherapy\|\| | 9 (60%) |
| Conventional systemic therapy<sup>+</sup> | 5 (33%) |
| Biologic agent | 9 (60%) |

PASI scores range from 0 to 72, with higher scores indicative of more severe disease. PGA evaluates psoriasis on a scale from 0 to 5, with 0 indicating cleared psoriasis, 1 minimal psoriasis, 2 mild psoriasis, 3 moderate psoriasis, 4 marked psoriasis, and 5 severe psoriasis.
\*Plus-minus values are the standard deviations from the mean
^Race was self-reported.
|| Phototherapy included ultraviolet B therapy and psoralen plus ultraviolet A therapy.
<sup>+</sup>Conventional systemic therapies include methotrexate, acitretin, and cyclosporine.

**Figure 1.**
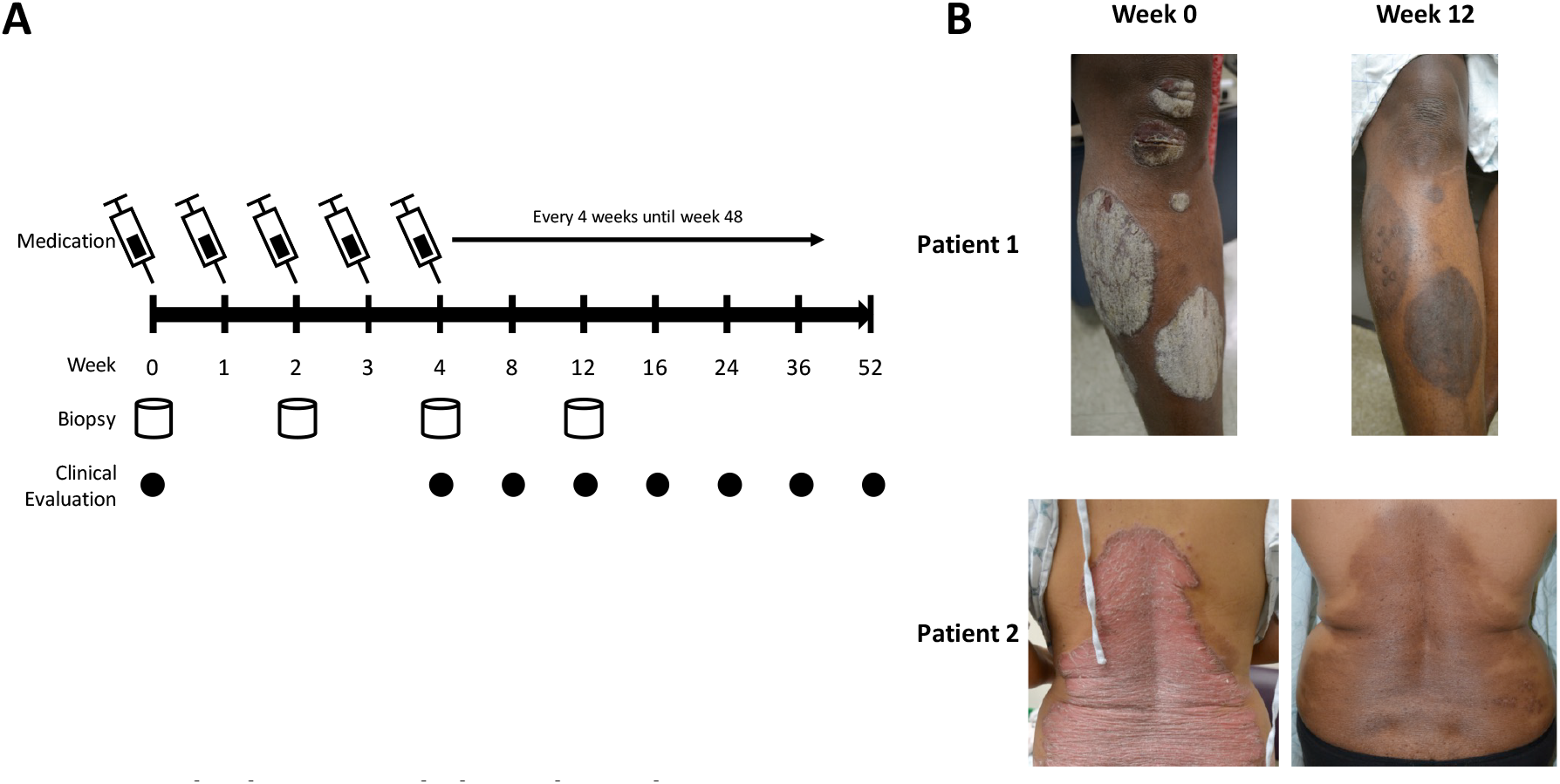
Study design and clinical results. (A) Study timeline. (B) Clinical improvement in two patients responding to secukinumab.

### Clinical improvement in response to secukinumab

Clinical measures of psoriasis severity improved in response to secukinumab. At week 12, plaques were substantially reduced (Figure 1B) and 100% of patients achieved PASI75. In terms of the speed of response, the cumulative mean reduction in PASI at weeks 4, 8, and 12 was 65%, 86%, and 90%, respectively. The mean PGA of patients was 4.0 at week 0 and 1.3 at week 12 (Table 2). PASI and PGA remained decreased until week 36 for most patients, with six patients showing an increase in PASI and PGA at week 52 during the period where secukinumab administration was not directly observed. Of the patients with reduced PASI at Week 52, three were observed to respond rapidly to treatment, achieving PASI100 at Week 8 and remaining at PASI100 for all subsequent evaluations.

**Table 2.** Average Clinical Response to Secukinumab

| Week | 0 | 4 | 8 | 12 | 16 | 24 | 36 | 52 |
| --- | --- | --- | --- | --- | --- | --- | --- | --- |
| PASI | 20.4 | 7.1 | 2.9 | 2.0 | 2.2 | 2.0 | 2.0 | 4.9 |
| PGA | 4.0 | 2.7 | 1.7 | 1.3 | 1.3 | 1.3 | 1.4 | 2.0 |

### Whole skin and CD8^+^ T cell-specific downregulation of IL-17 and IL-23 signaling in response to secukinumab

To investigate the transcriptomic response to secukinumab, we biopsied lesional skin from psoriatic patients before treatment and at 2, 4, and 12 weeks of treatment for comparison with skin biopsies from 13 healthy subjects. We performed RNA-seq on bulk skin tissue as well as on CD8^+^ T cells, Teffs, and Tregs sorted from these biopsies, sequencing each sample to an average depth of 41M reads per sample. Of the 271 collected samples, 241 (89%) passed QC, providing 9 – 14 samples per cell type per time point for analysis.

Within lesional skin, psoriasis-associated inflammatory gene expression was largely normalized during the course of secukinumab treatment. *IL17A*, *IL17F*, and *IL23A*, along with the genes encoding their major receptors *IL17RA, IL17RC, IL23R* and *IL12RB1* were upregulated in untreated lesional skin relative to healthy skin (Figure 2). These differences were either significant or trended toward significance (e.g. FDR-adjusted p-values of 0.059 and 0.085 for *IL17A* and *IL23R*, respectively). By week 12 of treatment, the expression of these genes had decreased to levels that were not significantly different from healthy samples. Disease-associated upregulation of *IFNG* was also reversed with treatment, but no significant disease-or treatment-related changes were observed in the expression of *TNF*, consistent with the modest changes observed in a previous study [8].

**Figure 2.**
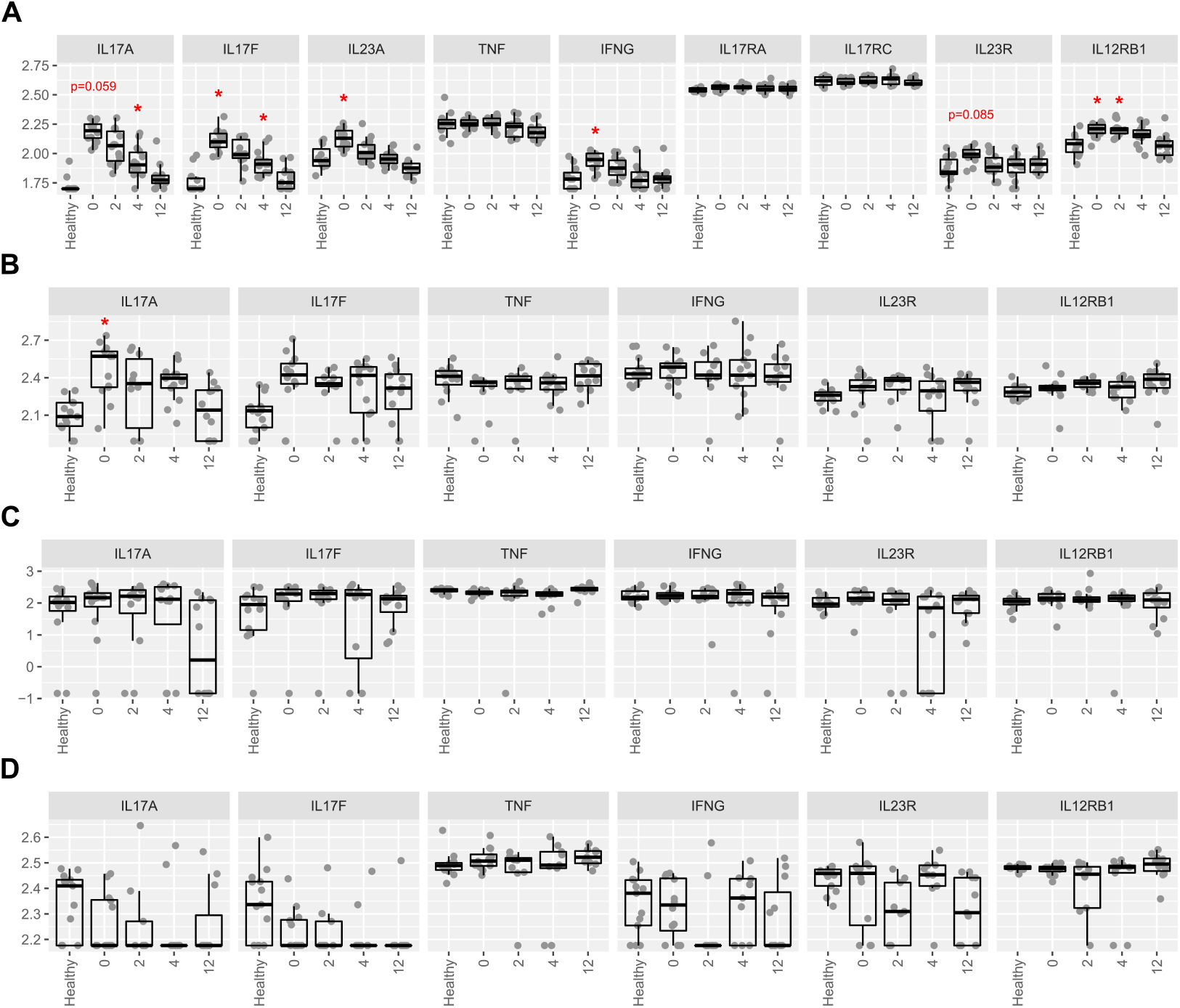
Expression of psoriasis-associated inflammatory genes is reduced in whole tissue and CD8^+^ T cells during secukinumab treatment. DESeq2-normalized and variance-stabilized expression of inflammatory genes in healthy, treated, and untreated samples for (A) bulk tissue, (B) CD8^+^ T cells, (C) Teffs, and (D) Tregs. Significance of differences based on Wald tests from separate DESeq2 models calculated between healthy samples and the samples of each timepoint. * = FDR-adjusted p-value < 0.05.

Among T cell subsets, only CD8^+^ T cells expressed significantly elevated *IL17A* expression in psoriatic conditions, and by week 12 of secukinumab treatment, the expresssion of this gene was again reduced to levels not statistically different from healthy samples. Psoriatic Teffs and Tregs, on the other hand, did not display abnormal differences in *IL17A, IL17F, TNF*, or *IFNG* levels, and expression of these genes in these subsets was not significantly altered by secukinumab.

### Global transcriptional profile of secukinumab-treated lesions and T cell subsets

A more general comparison of expression profiles revealed a broad shift in lesional gene expression over the course of secukinumab treatment. A PCA of 25,608 genes detected among the 66 bulk tissue samples showed a marked separation between 14 lesional samples biopsied before treatment and 11 healthy skin samples along the first and second principal components (Figure 3A). Lesional skin biopsied at subsequent weeks of treatment mark a progression toward the 11 healthy samples along PC1 but remain largely separate from the healthy samples, even by week 12 of treatment, when most patients had achieved PASI75.

**Figure 3.**
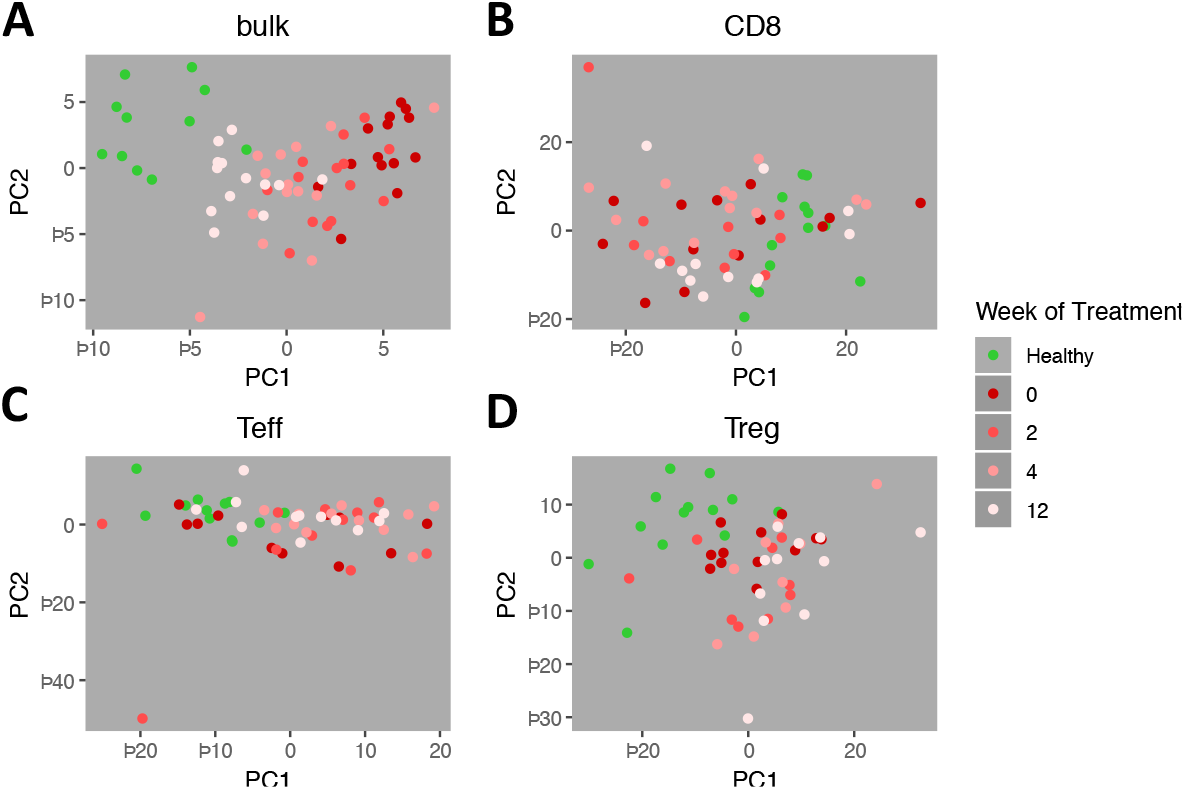
Broad shifts in lesional gene expression in response to secukinumab. PCA plots of (A) bulk tissue, (B) CD8^+^ T cells, (C) Teffs, and (D) Tregs. PCA was performed on normalized expression data from healthy, treated, and untreated samples of each sample type that were adjusted for the estimated effects of technical variables and variance-stabilized.

In contrast to the consistent shift in gene expression seen in bulk lesional tissue, we found more variability among sorted T cell populations in terms of their transcriptomic profiles and their response to secukinumab. In PCA plots of gene expression profiles from sorted CD8^+^ T cells, Teffs, and Tregs, we found some separation between healthy and lesional samples (Figure 3B-D) but no consistent progression during treatment.

### Disease-resolving and treatment-associated expression changes in response to secukinumab

We surveyed the transcriptomic changes that occur during secukinumab treatment by enumerating the differentially expressed (DE) genes between Week 0, Week 12, and healthy samples (Table 3, Supplementary Table 2). The number of DE genes detected between disease states and treatment conditions varied widely (11–3,096), which may reflect varying magnitudes of perturbation in psoriasis or response to treatment in bulk skin and different T cell subsets.

**Table 3.** Differentially expressed genes between healthy, untreated, and treated samples

| Sample type | Week 0 vs Healthy up/down | Week 0 vs Week 12 up/down | Week 12 vs Healthy up/down |
| --- | --- | --- | --- |
| bulk | 1289/1807 | 900/1165 | 96/186 |
| CD8<sup>+</sup> T cell | 40/89 | 2/26 | 7/18 |
| Teff | 9/28 | 3/11 | 1556/99 |
| Treg | 13/62 | 22/17 | 76/126 |

Most disease-associated expression differences in bulk tissue and sorted cells were resolved during secukinumab treatment. Of the 3,096 DE genes we found between healthy skin and untreated psoriatic lesions, 2,919 (94%) no longer showed significant differences between healthy skin and psoriatic lesions sampled at week 12 of treatment (Figure 4A). Similar resolution of disease-associated expression differences was found among the sorted T cell samples, with an average of 94% of DE genes in each subset restored at week 12. The general magnitude of expression differences in these DE genes was also progressively reduced throughout the first 12 weeks of treatment (Figure 4B), indicating reduced disease severity in response to secukinumab that is consistent with the drastic clinical improvement we observed above.

**Figure 4.**
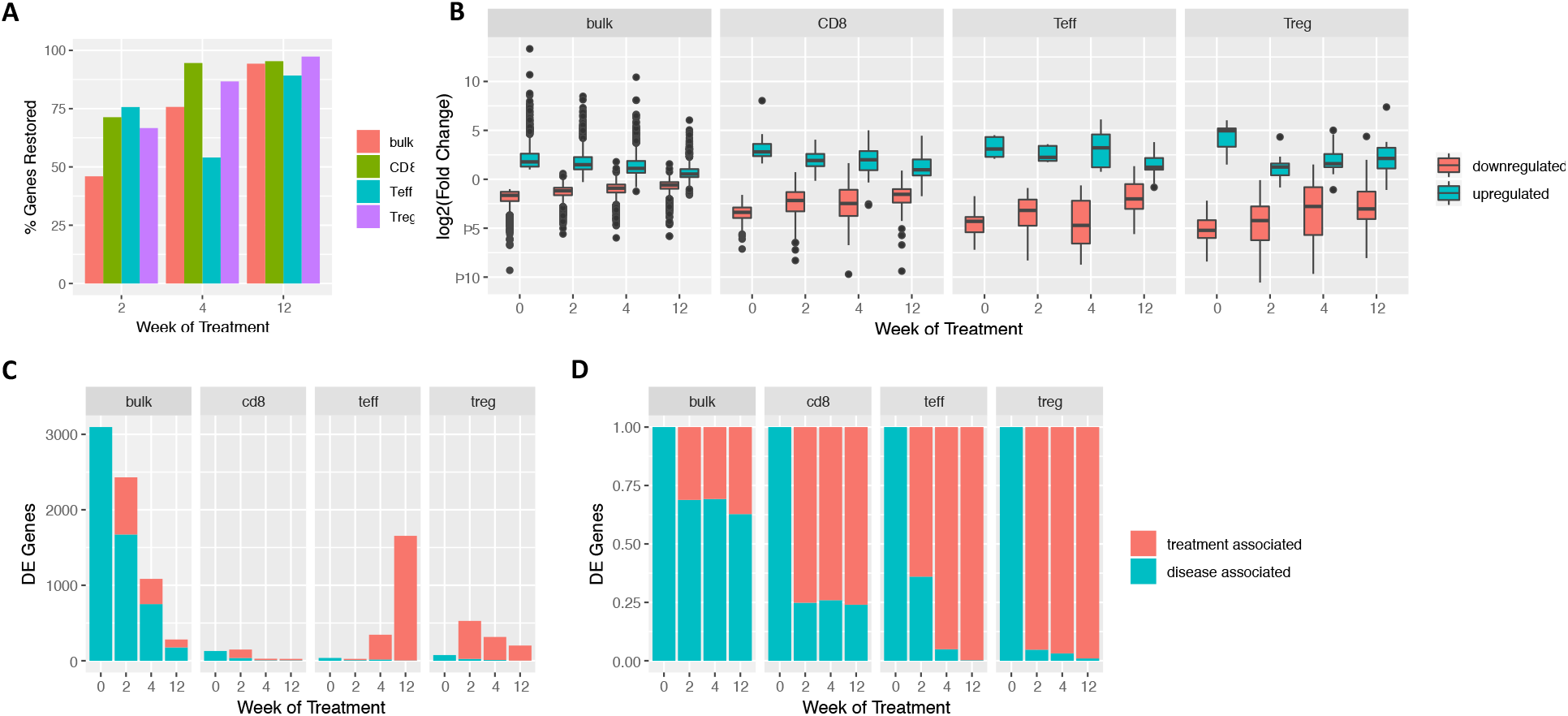
Secukinumab restores disease-associated transcriptomic changes and leads to treatment-associated expression differences. (A) Percentage of DE genes between healthy skin and untreated psoriatic lesion samples resolved at each timepoint of treatment. (B) log fold changes of these genes at each time point, which have been classified as up-or down-regulated based on their log2 fold change between psoriatic and healthy samples. (C) Number and (D) percentage of disease-associated or treatment-associated DE genes at each time point.

At the same time, secukinumab treatment also associated with, to varying extents, gene expression changes in bulk tissue and T cell subsets that we had not identified as DE between untreated psoriatic lesions and healthy skin. As summarized in Table 3, bulk tissue from psoriatic lesions at week 12 of treatment showed fewer significant expression differences from healthy skin (282 genes) compared to before treatment, and most of these (64%) were disease-associated expression differences that remained unresolved at this time point (Figure 4C, D).

On the other hand, DE genes between lesional and healthy CD8^+^ T cells, while also reduced in number during treatment, instead consisted mostly of genes not identified to be DE between untreated psoriasis and healthy conditions. Such “treatment-associated” genes compose a vast majority of the DE genes identified in Teffs and Tregs for each timepoint and increase in number within these subsets during the course of treatment (Figure 4C,D, Supplementary Table 2). Additional study is required to determine if these DE genes are due to treatment, the natural course of plaque resolution, or technical noise.

### Upregulated keratinocyte differentiation and immunological response pathways in psoriasis lesions compared to healthy skin

To gain insight into the biological processes underlying the transcriptomic differences between healthy, psoriatic, and treated skin, we used the Panther database [15] to identify pathways that are overrepresented in each set of DE genes. Comparison of DE genes between untreated psoriasis bulk skin and healthy bulk skin (week 0 vs. healthy, Figure 5A,C) revealed many pathways important for psoriasis pathogenesis. Psoriasis skin has higher expression of genes involved in keratinocyte differentiation (FDR=0.01), epidermis development (FDR=0.003), and the mitotic cell cycle process (FDR<10^19^), consistent with a hyper-proliferation of keratinocytes that are characteristic of psoriasis. Neutrophil activation genes (FDR=0.0002) are also elevated in lesional samples, reflecting increased neutrophil infiltration and activation in psoriatic skin. Lastly, DE genes upregulated in psoriasis skin are also enriched with interferon gamma (FDR=0.0001), IL-17 (FDR=0.003), and IL-23-signaling pathways (FDR=0.02)

**Figure 5.**
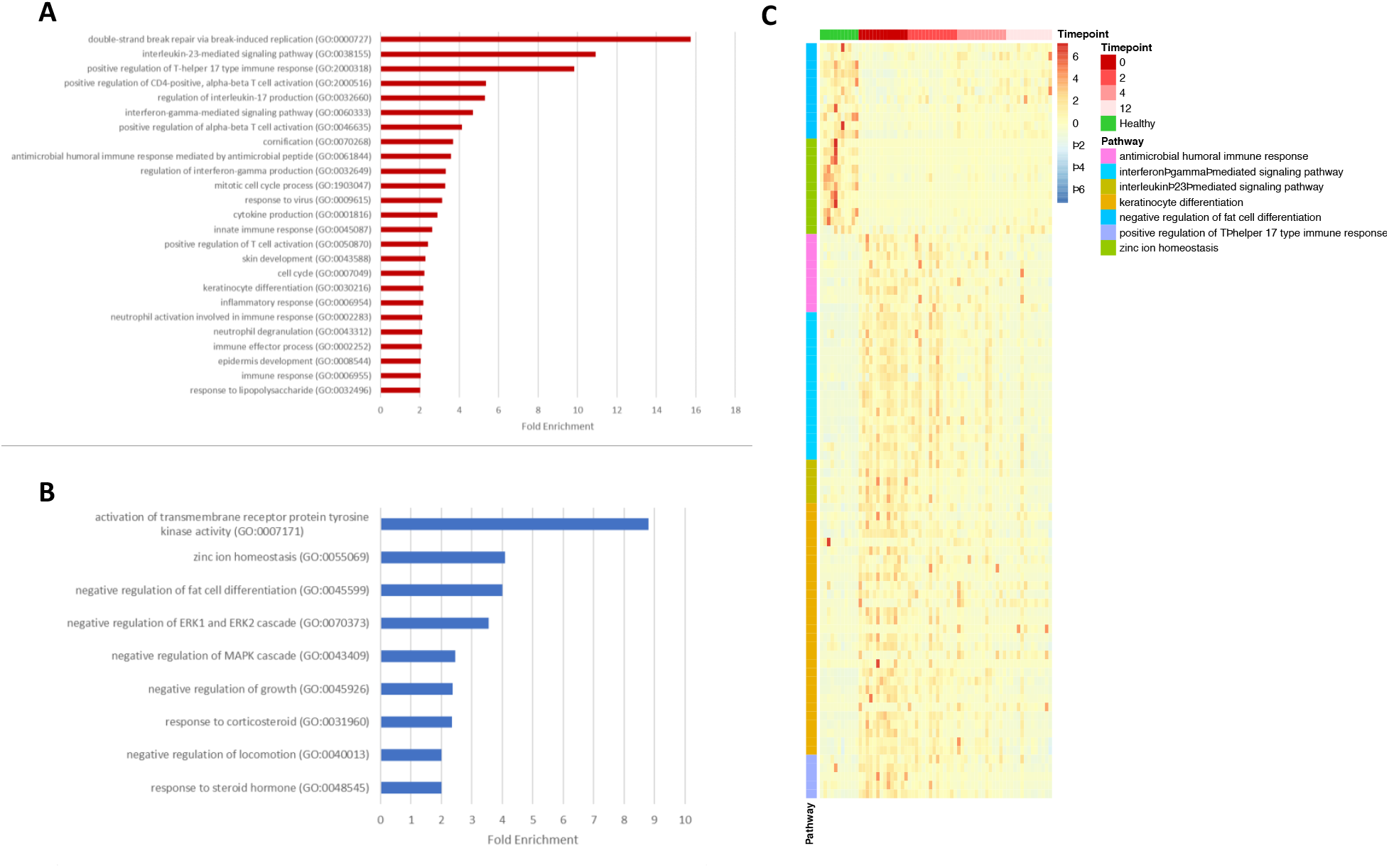
Pathways overrepresented between psoriatic and healthy skin. Gene Ontology pathways overrepresented among the (A) upregulated and (B) downregulated DE genes between psoriatic and healthy bulk tissue. (C) Normalized expression of genes in each pathway across treatment timepoints.

On the other hand, the genes that are down-regulated in psoriasis skin compared to healthy skin (Figure 5B, C) contain an overrepresentation of genes involved in negative regulation of locomotion (FDR=0.01), MAPK signaling (FDR=0.01), and ERK signaling (FDR=0.007), consistent with increased migration and activation of T cells commonly observed in psoriatic skin. Down-regulated genes were also enriched with genes involved in negative regulation of fat cell differentiation (FDR=0.02) and genes involved in zinc ion homeostasis (FDR=0.04). Many of the ion homeostasis genes encode metallothioneins, which are small, ubiquitous, cysteine-rich proteins with antioxidant properties that have been implicated in mediating anti-inflammatory effects in various mouse models [16]. No biological process or pathway was found to be overrepresented for psoriatic CD8^+^ T cells, Teffs, and Tregs when compared to their healthy counterparts or across treatment timepoints, as only a modest number of DE genes were identified for these subsets.

## Discussion

In our study, secukinumab treatment was associated with clinical improvement as well as a dampening of disease-relevant gene expression and pathways in whole lesional skin, consistent with previous studies [8,9]. We additionally observed that T cell subsets in secukinumab-treated lesions showed a similar resolution of expression in genes perturbed by disease. However, the transcriptomic differences between T cell populations in healthy and psoriatic skin as well as in psoriatic skin before and after treatment were limited, despite well documented evidence of an activated phenotype acquired by T cells in psoriasis [10]. This may be explained by the higher degree of inter-subject variability we observed within these cell subsets relative to bulk tissue, the chronicity of disease perhaps leading to T cells in a non-activated equilibrium state, as well as the possibility that the diseased-involved or secukinumab-responsive cells may make up only a small proportion of the total number of cells of each subset in the lesion.

Further comparison of T cells in healthy skin and psoriatic lesions as well as pre- and post-treatment revealed other observations. While it is generally understood that both CD4^+^ and CD8^+^ T cells can produce IL-17 in psoriatic lesions [4,17] and respond to autoantigens [10], in our study *IL17A* overexpression predominates in psoriatic CD8^+^ T cells rather than CD4^+^ T cells, which may reflect greater importance of Tc17 compared to Th17 cells as sources of IL-17A.

Additionally, we observed treatment-associated gene expression changes in both bulk tissue as well as T cell subsets that significantly differed from their healthy counterparts during secukinumab treatment, which is consistent with our PCA analysis suggesting that the expression profiles of bulk tissue and sorted T cells in secukinumab-treated lesions remain distinct from that of healthy skin. These expression differences may reflect a number of possible scenarios, such as genes directly altered by treatment, the activation of additional genes and pathways that may be involved in lesion resolution, as well as a distinct “treatment-resolved” state, as has been described for etanercept treatment [18], where resolved skin was observed to harbor a persistent population of Tc17 clones. These observations warrant further investigation using higher-resolution technologies, such as single cell RNA-seq, for profiling the expression and phenotypes of lesional T cell subsets.

## Acknowledgments

The authors would like to thank Mary Patricia Smith and Karen Ly for their assistance with this project.

## Supplementary Information

Supplementary Table 1. Patient demographics.

Supplementary Table 2. Differentially expressed genes between samples of each treatment timepoint and healthy samples.

